# Indigenous Knowledge as a sole data source in habitat selection functions

**DOI:** 10.1101/2023.09.07.556613

**Authors:** Rowenna Gryba, Andrew VonDuyke, Henry Huntington, Billy Adams, Brower Frantz, Justin Gatten, Qaiyyan Harcharek, Robert Sarren, Greg Henry, Marie Auger-Méthé

## Abstract

While Indigenous Knowledge (IK) contains a wealth of information on the behaviour and habitat use of species, it is rarely included in the species-habitat models frequently used by ‘Western’ species management authorities. As decisions from these authorities can limit access to species that are important culturally and for subsistence, exclusion of IK in conservation and management frameworks can negatively impact both species and Indigenous communities. In partnership with Iñupiat hunters, we developed methods to statistically characterize IK of species-habitat relationships and developed models that rely solely on IK to identify species habitat use and important areas. We provide methods for different types of IK documentation and for dynamic habitat types (e.g., ice concentration). We apply the method to ringed seals (natchiq in Iñupiaq) in Alaskan waters, a stock for which the designated critical habitat has been debated in part due to minimal inclusion of IK. Our work demonstrates how IK of species-habitat relationships, with the inclusion of dynamic habitat types, expands on existing mapping approaches and provides another method to identify species habitat use and important areas. The results of this work provide a straightforward and meaningful approach to include IK in species management, especially through co-management processes.

“Agencies have a traditional way they do science and including Indigenous Knowledge is less traditional.” - Taqulik Hepa, subsistence hunter and Director, North Slope Borough Department of Wildlife Management

**Statement of Positionality:** This study and the conversion and application of Indigenous Knowledge (IK) for habitat use models was initiated through discussions with the North Slope Borough Department of Wildlife Management (DWM). The DWM is an agency of the regional municipal government representing eight primarily Iñupiat subsistence communities in Northern Alaska. One of the goals of the DWM is to “assure participation by Borough residents in the management of wildlife and fish… so that residents can continue to practice traditional methods of subsistence harvest of wildlife resources in perpetuity” (1). Additionally, this project was presented to the Ice Seal Committee (ISC) for review, input, and approval. The ISC is an Alaskan Native organization with representatives from five regions that cover ice-associated seal ranges and “was established to help preserve and enhance ice seal habitat; protect and enhance Alaska Native culture, traditions-particularly activities associated with the subsistence use of ice seals” (2). Both the DWM and the ISC have mandates to manage ice-associated seals considering both IK and ‘Western’ scientific knowledge (1, 2), and this study was developed to meet those mandates. Iñupiat hunters from Utqiaġvik, Alaska (Figure 1) were collaborators on this project, five of whom are co-authors (B. Adams, B. Frantz, J. Gatten, Q. Harcharek, and R. Sarren), while the other hunter chose to remain anonymous for this publication. The other authors are not Indigenous: R. Gryba was a PhD candidate at the University of British Columbia, M. Auger-Méthé and G. Henry are professors at the University of British Columbia, A. Von Duyke is a researcher at the DWM, and H. Huntington is an independent social scientist.

**Significance Statement:** Indigenous Knowledge (IK) is an extensive source of information of species habitat use and behavior, but is still rarely included in statistical methods used for species conservation and management. Because current conservation practices are frequently still rooted in ‘Western’ practices many Indigenous organizations are looking for ways for IK to be better included and considered. We worked with Iñupiat hunters to develop a new statistical approach to characterize IK and use it as a sole data source in habitat models. This work expands on mapping approaches, that are valuable, but cannot be applied to dynamic habitat types (e.g., ice concentration). This work shows how IK can be meaningfully included in modelling and be considered in current approaches for species management.

## Introduction

Current species conservation and management practices frequently lie within a ‘Western’ science context (3–5). This focus on ‘Western’ science and management practices often leads to superficial or a lack of consultation with Indigenous Peoples (6, 7), and limited inclusion of, and respect for, IK (7–9). Exclusion of Indigenous management practices and knowledge, and the focused application of ‘Western’ scientific management practices has led to, for example, poor fisheries management (10) and hunting bans that can impact the health, well-being, and livelihoods of Indigenous communities (11). The recognition that ‘Western’ species management has often failed to achieve its goals has brought about necessary discourse on shifting species management to center IK and Indigenous management frameworks, led by Indigenous Peoples (12–15). This discord has lead to increased efforts to recognize the value of IK and IK systems in conservation management (16–18), recognition of the importance of knowledge co-production (19–21), co-management (22–24), and Indigenous-led conservation (25). When IK is valued and included in conservation and management decision making the decisions are based on a broadly informed understanding of species that includes multiple ways of knowing (e.g., 12,20,24).

**Fig. 1.**
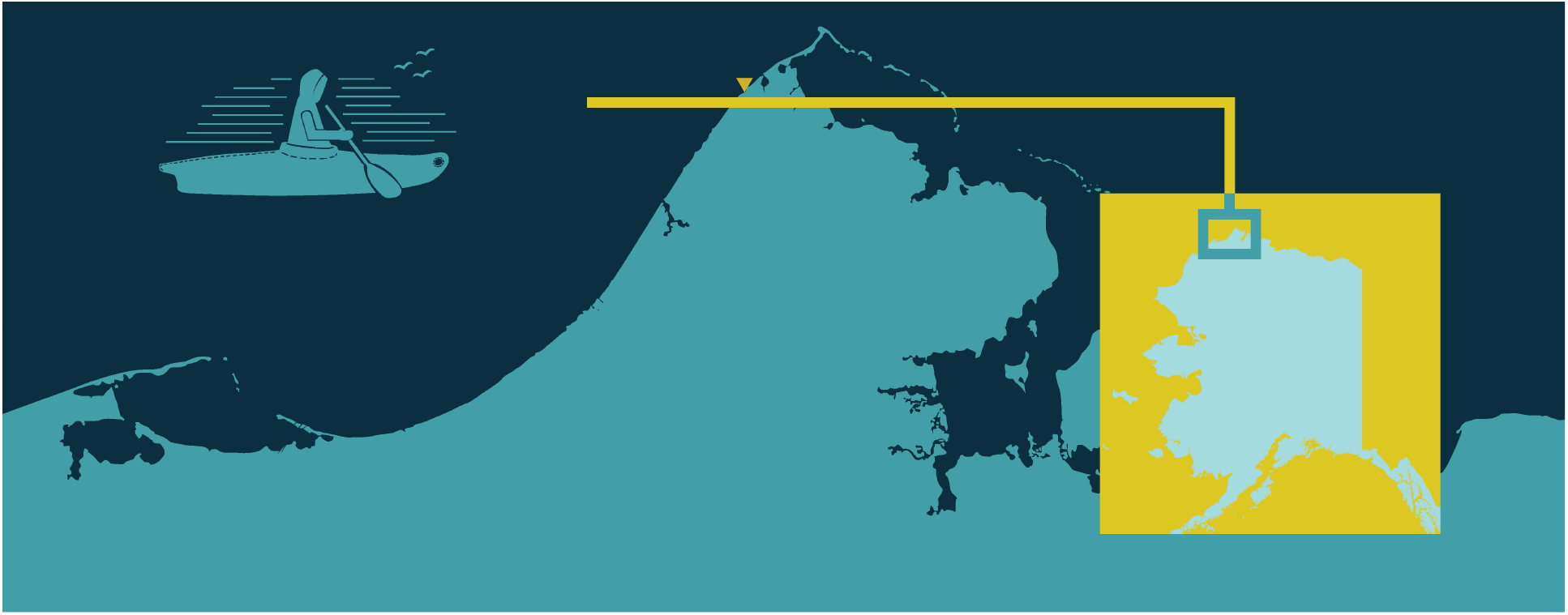
Study area around Utqiaġvik, Alaska (shown in the yellow triangle).

Although increases in the inclusion of IK in conservation practices are encouraging (18), ‘Western’ science species conservation management often relies on characterizing habitat use and identifying critical habitat using statistical tools such as resource selection or habitat selection functions (HSFs) that rely on ‘Western’ science data alone (26). These tools are used to estimate how the occurrence of a species varies across space, and are developed from animal occupancy or movement data (e.g., satellite telemetry data). Habitat suitability analysis has informed 40% of critical habitat definitions for Canadian vertebrate species at risk, providing information on species-habitat relationships (27). Indigenous Knowledge contains, among many other things, information on species habitat use and can be used to inform the identification of critical habitat. Indigenous Knowledge can be defined as “Knowledge created and/or mobilized by Indigenous Peoples that may include Traditional Knowledge and scientific knowledge.” (28, pg. 719). As noted by co-author and Iñupiat hunter B. Adams, knowledge gained by hunters over time can have a better understanding of species habitat use and behavior than approaches that just use satellite tagged animals.

The formal combination of IK and ‘Western’ scientific methods has been limited (29) but is increasing in ecological modelling (30, 31). Previous efforts to include IK in ‘Western’ scientific approaches often relied on comparison between the two knowledge systems (e.g., 32–34). The reasons for this comparative approach are varied but include how IK is perceived (e.g., 13, 17) and the lack of statistical methods for more formal inclusion of IK and ‘Western’ scientific data in the same approach. IK has been included in previous studies of habitat use (35–37), but those studies do not directly characterize or estimate the relationships between species and habitat covariates. Expanding on these methods to be able to directly estimate IK species-habitat relationships and the ability to consider changing habitat due to climate change will provide additional approaches to better inform species management and conservation, further recognizing IK as a valuable knowledge source.

One of the reasons for the increased inclusion of IK in species management is the increase in co-management agreements between Indigenous governments/organizations and non-Indigenous governments. In both Canada and the US, Indigenous co-management organizations represent local or regional priorities, and are linked to local management organizations (e.g., Hunter Trapper Associations in Canada, North Slope Borough Department of Wildlife Management in Alaska, US) and to federal agencies (e.g., Fisheries and Oceans Canada, National Marine Fisheries Service). Although several co-management frameworks are in place in many regions (e.g., 22), most often the final decision on species listings and critical habitat designations still sit with federal agencies. For example, in the United States, the ISC has a co-management agreement in place with the National Marine Fisheries Service (NMFS) (38), but final decisions regarding the listing and critical habitat designation of ice-associated seals still lie with NMFS (39). Although the best solution would be the full consideration of Indigenous management and understanding on its own terms (9), current co-management practices often create the need to convert IK into a form that fits both ‘Western’ and Indigenous management methods (24, 40). The method we propose in this paper is developed within this context and we acknowledge that efforts to convert IK into a form that can be included in ‘Western’ scientific methods can be extractive, separating IK from its multi-dimensional context (6, 41). However, the research was developed following a knowledge co-production framework (19) and to meet the needs of Indigenous partners. It reflects the desire for IK to be more readily included into ‘Western’ science approaches for conservation management and the need for IK to be fully considered in co-management agreements. Here, we develop a framework that uses IK as sole source of information (i.e., no other species observation data) to statistically estimate species habitat selection, including dynamic habitat types. The framework uses a two-step approach inspired by one used in classic habitat selection analyses (Figure 2). The first step involves using IK to characterize the species relationship with each environmental covariate, considering different ways that IK may be documented. Knowledge documented via semi-directed interviews comes in many forms (e.g., maps of where animals are observed, types of habitat animals are associated with; Figure 3A, B), which affects how the relationship with a given covariate is characterized. As such, we classify the IK documentation into three types (Figue 3A,B): quantitative (e.g., species are associated with a specific habitat value), qualitative (e.g., category of habitat animals are associated with), and spatial (e.g., maps of where animals are observed). Each IK documentation type is characterized through different means (e.g., beta regression for quantitative; Figure 3, see methods for detail). The second step involves combining the relationship with all covariates to predict the overall probability of use. The IK holders review the results at both steps (Figure 2).

**Fig. 2.**
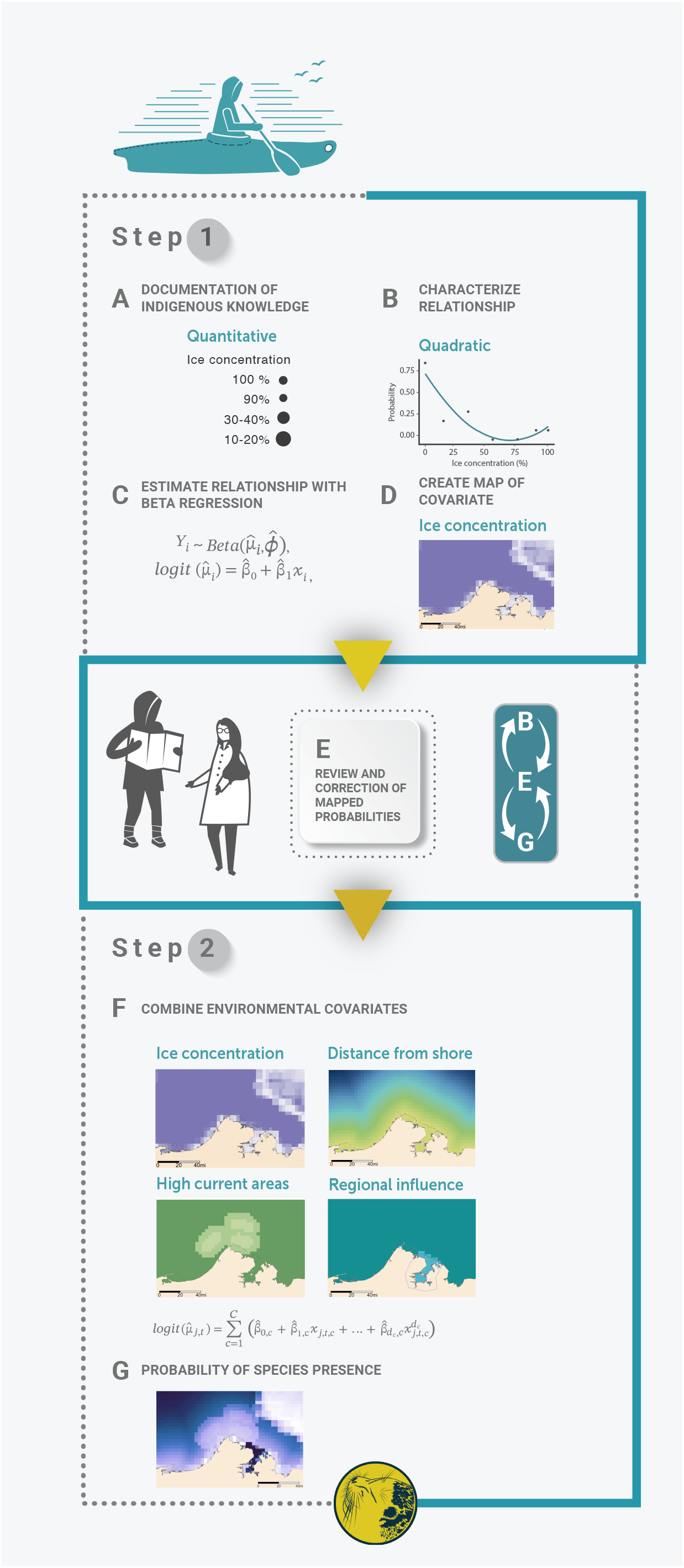
Methods process map of (A) the documentation of IK, (B) characterization of relationship between IK probability of species presence and habitat covariates, (C) application of beta regression to quantitative covariates, (D) mapping of IK probability of presence for review, (E) review and correction of the IK probabilities of presence, (F) using *β* estimates in a logistic regression equation to estimate (G) the final probability of species presence, and reviewing the final estimate of probability of species presence for any needed corrections. The size of the circles in (A) represent the proportion of hunters that indicated that species-habitat association and inform the probability values in (B).

**Fig. 3.**
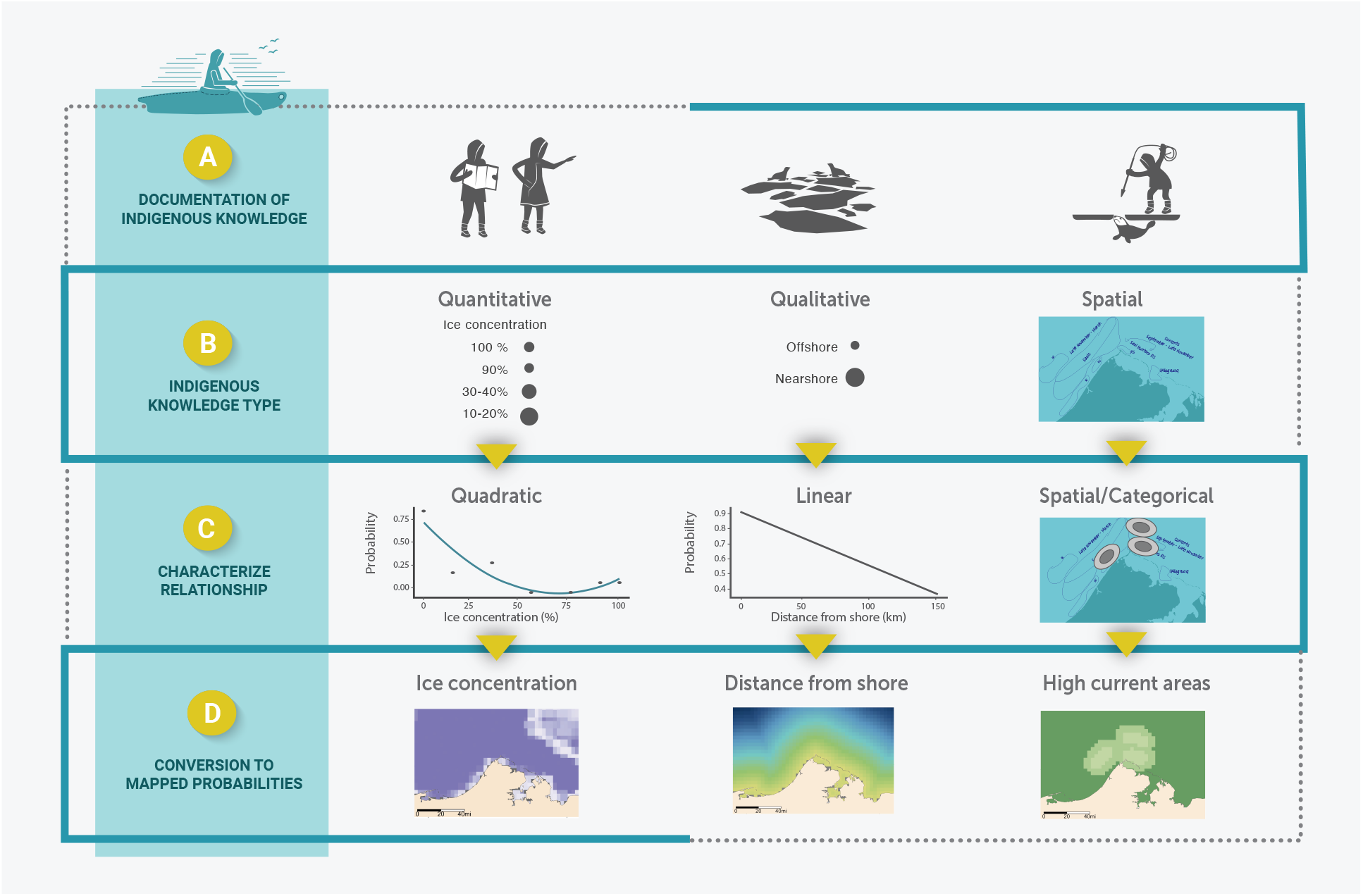
Methods process map of (A) the documentation of Indigneous Knowledge (IK), (B) identification of IK type, (C) characterization of relationship between IK probability of species presence and habitat covariates, (D) mapping of IK probability of species presence. The size of the circles in (B) represent the proportion of hunters that indicated that species-habitat association and inform the probability values in (C).

Our new method can account for dynamic habitat types (e.g., sea ice concentration) and complex species-habitat relationships, that would be difficult to model using ‘Western’ scientific data alone (e.g., appropriate data may not exist, data is at a too coarse spatial resolution). Our method can be used to map spatial variation in species presence, identify important areas, and creates estimates of species habitat use that fully consider IK. To demonstrate our approach we apply the methods to a case study on ringed seals (*natchiq* in Iñupiaq; *Pusa hispida*). Ringed seals are an important species culturally and for subsistence for many Inuit (42, 43) and have recently been designated under the US Endangered Species Act (44, 45).

## Results

We applied the methods to four environmental covariates that IK holders identified as influencing the habitat selection of ringed seals in summer in the waters around Utqiaġvik, Alaska (Figure 1): distance from shore, ice concentration, currents, and spatial regions. The relationships between ringed seals and the environmental covariates defined here were based on IK documented in Gryba *et al*.(46), although some adjustments were made during review of the maps with the hunters. Knowledge of distance from shore, a qualitative type of IK, indicated a higher probability of ringed seal association with nearshore compared to offshore habitat (46), which we characterized as a simple linear relationship (Figure 4A, Table 1). During the review of the probability of ringed seal presence maps, one hunter noted that the region of Admiralty Bay should have lower probability of presence, a spatial type of IK. The probability of presence within the specified region was decreased accordingly (Figure 4B, Table 1). Ice concentration was treated as a quantitative type of IK, and ringed seal presence was quantified for a set of ice concentration bins (i.e.,0%, 10-20%, 30-40%, 50-60%, 70-80%,

**Table 1.** IK based coefficient values (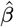 and *σ*^2^) for each habitat type.

| Model | Habitat type | $\hat{\beta}$ | $\sigma$ |
| --- | --- | --- | --- |
| Quantitative | Intercept | 0.12 | 0.86 |
|  | Ice concentration | 1.51 | 0.50 |
|  | Ice concentration<sup>2</sup> | 4.48 | 0.77 |
| Qualitative | At shore (intercept) | 0.88 | — |
|  | Distance from shore | -0.16 | — |
| Spatial | Current regions - outer (intercept) | 0 | — |
|  | Current regions - near (10-20 km) | 0.08 | — |
|  | Current regions - main | 0.11 | — |
|  | Admiralty Bay region - inside (intercept) | 0 | — |
|  | Admiralty Bay region - outside | 0.89 | — |

**Fig. 4.**
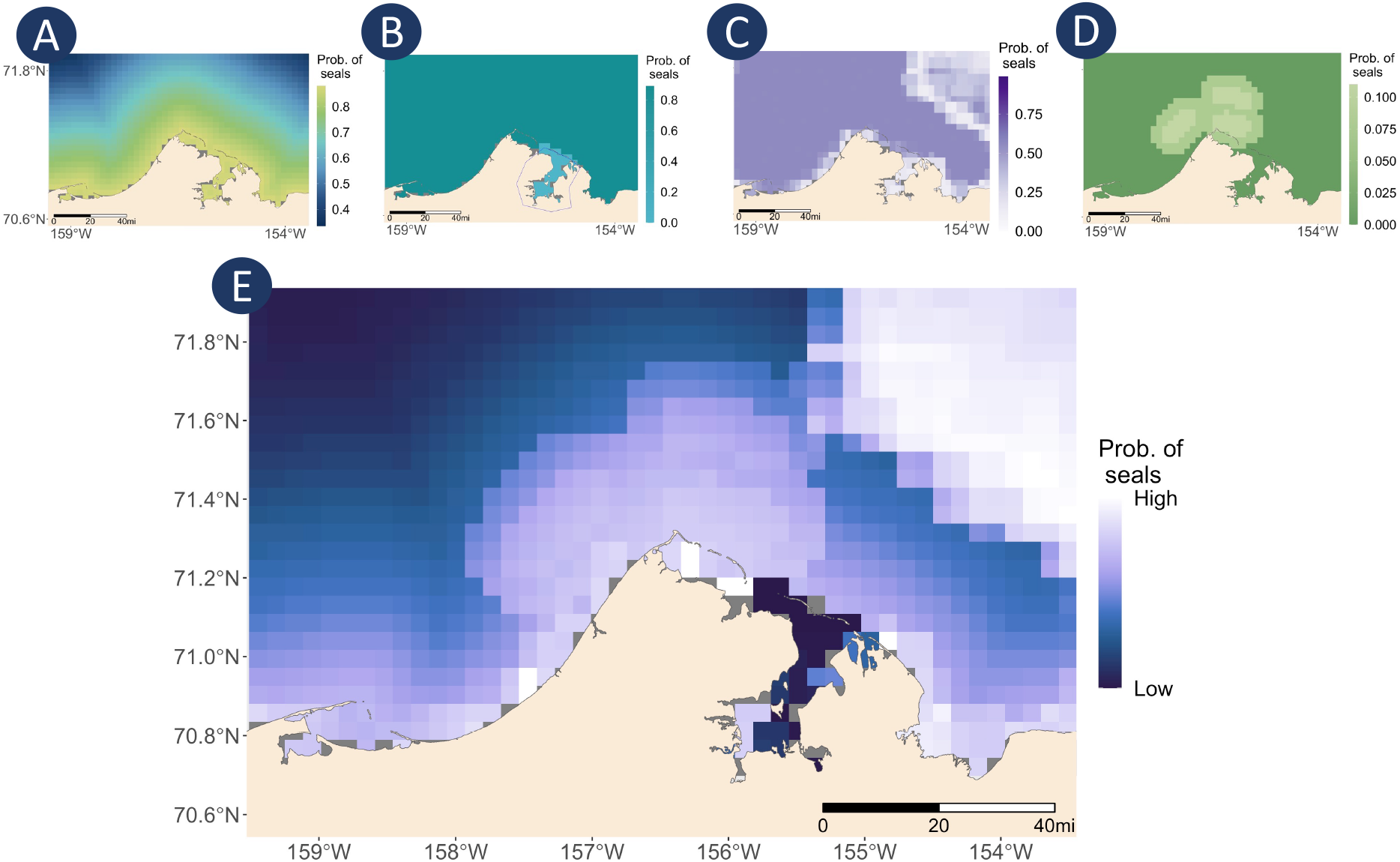
IK probability of ringed seal presence associated with habitat covariates and final predicted habitat selection. (A) IK probability of ringed seals presence associated with distance from shore. Yellow indicates high probability of presence and blue indicates low probability of presence. (B) IK probability of ringed seal presence associated with distance from shore with decreased probability within Admiralty Bay. Dark blue indicates high probability of presence areas, light blue indicates low probability of presence, Admiralty Bay is enclosed by the polygon shown in blue. (C) IK probability of ringed seal presence associated with ice concentration for July 15, 2015. White indicates low probability of presence and dark purple indicates high probability of presence. (D) IK probability of ringed seal presence associated with currents. Light green indicates areas of higher probability of presence, dark green indicates areas of lower probability of presence. (E) Final IK probability of ringed seal presence for July 15, 2015. White indicates areas of high probability of presence, dark blue indicates ares of low probability of presence.

90%, 100%) (46). While reviewing the IK probability maps that included a large area of ice (i.e., July 2015, Figure S1), the knowledge holders indicated that numerous seals would be present in areas of high ice concentrations. As such, the probabilities in high ice concentration were increased (Figure 4C). The knowledge holders indicated that these increases better reflected their observations. We used a quadratic beta regression to quantify this relationship (Table 1). Hunters indicated areas of currents used by ringed seals in summer (46), a spatial type of IK. The identified areas were given a probability of presence of 0.11 reflecting that each region was identified by one hunter (i.e., 1 hunter out of 9 hunters). Areas 10-20 km around these regions were given probabilities of 0.08 to provide a buffer around the currents (Figure 4D, Table 1). The seal habitat use was predicted by combining into a single equation (using equation 5 in the Methods) all of the variables and the estimated coefficients characterizing their relationship with seal presence (Figure 4E).

## Discussion

“Showing IK in this way is a useful step to get buy-in and show the value of IK for agencies, and for them to see the wealth and depth of IK. It shows the value of IK and the importance of IK as a data set.” - Taqulik Hepa, subsistence hunter and Director, North Slope Borough Department of Wildlife Management

Our method provides an approach to convert quantitative, spatial, and qualitative documentation of IK into probabilities which can subsequently be modelled and used to estimate habitat coefficients and species habitat use with IK as a sole data source. Our application of beta regressions in this context leans towards elicitation methods associated with ‘expert knowledge’ (47, 48). However, while IK can be classified as expert knowledge it, embodies a far greater depth of information about species habitat and is often documented using more semi-directed methods. Beta regression provides an opportunity to directly model proportions with clear applications to ecology (49, 50), and in our case an approach to model IK probabilities of presence to quantify species-habitat relationships. This approach, which can be applied to dynamic habitat types, is of particular importance in the context of climate change. Sea ice thickness and the timing of formation in the fall and retreat in the spring have been changing with climate change (51), and these changes have the potential to impact ringed, bearded, and spotted seals habitat use and distribution in some areas (e.g., 52–54). The methods here can be applied to predict changes in habitat use for dynamic habitat types, recognizing that the IK of ringed, bearded, and spotted seal habitat use may change over time reflecting any observed changes in habitat use.

While the implicit assumptions often associated with habitat modelling (e.g., sampling bias, model extents) (55) apply to our method, some are mitigated by the use of IK. For example, sampling bias frequently present in scientific studies (e.g., small sample size per year)(56, 57) is less present in IK. Hunters are observing multiple species throughout the year, which is reflected in the holistic nature of IK. The breath of knowledge and observations, and general understanding of common species, additionally address the assumption that data sampled for habitat models are representative of the population (55). There may be bias in the IK towards areas that are visited more frequently. We addressed this bias by restricting the model predictions to areas used by hunters (58) and would not extend the IK applied in our case study to other regions without prior verification with knowledge holders (e.g., assumptions regarding interpolation). Gryba *et al*. (46) highlights the importance of speaking to the Indigenous People of a place and that IK from different areas can reflect differences in species behaviour and habitat use. We do assume that the variables selected, and identified by the IK holders, reflect the appropriate systems, sometimes as proxy of what is influencing ringed seal habitat use. As in most habitat selection methods (e.g., 59, 60), our modelling approach assumes that the effects of environmental variables are additive. There are examples where this assumption does not hold (e.g., 61), but the review by the IK holders of the final output and indication that it reflects their IK leads us to be confident that this assumption is reasonable in this case. One of the main motivations for this study was to create estimates of species habitat use that fully consider IK and methods that can be applied to dynamic habitat types such as ice concentration. This was partially in response to the listing of ringed seals and bearded seals as threatened under the US Endangered Species Act, and subsequent designation of critical habitats (44, 45). The use of IK as a data source in critical habitat predictions provides an opportunity to reduce scientific uncertainty or lack of data for species that are data-limited in terms of ‘Western’ science approaches (62, 63). For example, satellite telemetry data of ringed seals is limited in summer in the waters near Utqiaġvik, Alaska as most individuals tagged move farther offshore, west into the northern Chukchi Sea, or east into the eastern Beaufort Sea (57). As such, telemetry data may not capture the habitat use of the ringed seals that remain in waters near Utqiaġvik. In contrast, IK provides information on the seals that remain in the region and is not limited by the small sample size associated with many telemetry studies. Additionally, covariates such as current can act as a proxy for foraging areas, given the association noted by IK holders (46). The ringed seal critical habitat designation is based on essential features that include sea ice suitable for whelping, nursing, and basking in waters deeper then 3 m (44), information that is primarily based on satellite telemetry and aerial survey data. The critical habitat was also assessed to sufficiently represent areas of ringed seal primary prey. The habitat use identified in this study indicates high use of ringed seals in nearshore habitat that is not included in the current critical habitat designation. Although the critical habitat is not intended to include all areas of ringed seal presence, consideration of IK on ringed seal use of the waters near Utqiaġvik, Alaska may reflect information not available in ‘Western’ scientific data sources. Additionally, the IK not only highlights areas of high use but also identifies areas important for foraging (46).

The quantification of IK in this study can provide an approach for IK to be a sole data source in habitat use modelling. These methods can also be used as a starting point to combine IK and ‘Western’ scientific data in the same models, using a Bayesian framework for example. Regardless, our approach inherently considers a small portion of what IK embodies as the links between species, people, and place are not included (64). Although only a portion of IK is considered in these types of methods, the approach can provide opportunities for co-management bodies to clearly show the extent of species-habitat relationships that are part of IK and that standard scientific data need not be the only data source considered in habitat modelling approaches. Our approach matches the methods often used with scientific data and allows for straightforward mapping of species relationship with dynamic habitats, simplifying the inclusions of results in current ‘Western’ science based management schemes. The efforts of co-management organizations to center IK in management can move forward the decolonization of current management authorities (3). This is particularly important considering the relationships that Indigenous Peoples have with species (28, 64) and the need to consider cultural and traditional practices and move forward with true reconciliation and restoration (3).

Translation of IK into forms such as habitat use may serve to provide opportunities for IK to be considered in new ways within many of the current dominant conservation management frameworks. The approach, if initiated and guided by IK holders and Indigenous communities, can provide opportunities for knowledge co-production that can help to communicate the extent of IK and also the relationships between Indigenous Peoples, their land, and species (3, 19). Additionally, the recognition and inclusion of IK as an integral data source can help solve many of the issues we are facing when attempting to manage species in a changing world (10, 19).

“One isn’t better than the other [IK and scientific data] and both are equally valuable and can give a broader understanding. This approach helps to show the depth of understanding in IK and there is a need to help grow the understanding of IK for management use.” - Taqulik Hepa, subsistence hunter and Director, North Slope Borough Department of Wildlife Management

## Materials and Methods

Habitat selection functions often have two sequential goals: 1) quantifying the relationship between a species presence and environmental covariates, and 2) using these estimated relationships to predict how the species occurrence vary spatially (60). Here, we use a similar two-step approach. We first use IK to characterize the relationship with each covariate, and then combine these relationships to create maps that display the spatially-varying probability of a species presence.

### Step 1: Using IK to characterize the species relationship with each environmental covariate

The first step of the approach requires quantifying the relationship between the probability of species presence and each environmental covariate. To quantify these relationships using IK, we assume that the proportion of knowledge holders interviewed who share information specific to a habitat covariate value is equivalent to the probability of species presence associated with that covariate value. For example, if only one of the *n* hunters interviewed declare that seals are present in areas with currents, the probability of seal being present at those areas is set to 1*/n*. This assumption aligns with the feedback given during semi-directed interviews in Gryba *et al*.(46), where hunters acknowledged that others have different experience and that less common behaviours may not be observed by all. This assumption also allows for the inclusion of all IK shared, and does not imply uncertainty or reduced confidence to less frequently observed behaviours or habitat use.

For the quantitative type of IK, the relationship between species presence and a covariate can be estimated using beta regression (Figure 42A-C). Beta regression is a distributional regression that can be used to model a continuous response variable *Y* within the interval (0,1), such as probability of presence or proportion (49, 65). A beta regression model can be defined in several ways, we used the following parametrization:

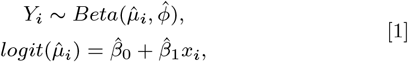

where *Y*_*i*_ is probability of species presence for a specific covariate value *i*, 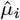 is the estimated expected mean value of *Y*_*i*_ associated with the value of the covariate, and 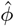 is the precision, *x*_*i*_ is the covariate value, and 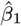 is the estimated regression coefficient that characterizes the relationship between the covariate and the IK-informed probabilities of species presence *Y*_*i*_, and 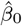 is the estimated intercept. The two *β* coefficients are estimated, and are the main parameters of interest in this step of our approach. Beta regression can also be used to characterize quadratic relationships.

Characterizing the qualitative and spatial types of IK (Figure 3C,D) does not require the Beta regression. Instead, the relationships are set based on the values of the covariate and the IK probability of species presence. For example, a linear relationship can be characterized by simply fitting a line and then calculating the slope and intercept of that line to define the relationship. Spatial IK (i.e., IK that was drawn on maps) can have regions that are higher or lower probability of species presence, based on a background value, with buffers to indicate change in probability of species presence (Figure 3C,D). The background value is the intercept (*β*_0_) and is assumed to be 0, with the other *β* coefficients measuring the difference from the background value. If needed, a transformation can be applied to ensure that the probabilities remain between 0 and 1.

Once the *β* coefficients are estimated, they are used to predict the probability of species presence to a rasterized version of the covariate and mapped (Figure 2D). All of the maps are reviewed by the knowledge holders originally interviewed and can be edited as needed (Figure 2E).

### Step 2: Combining the relationship with covariates to predict overall probability of use

The second step of the approach combines the relationship with each covariate into a single model that is used to predict the overall spatially-varying probability of a seal presence.

Simply, we can predict the probability of species presence in each cell *j* of our rasterized map using

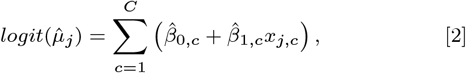

where *C* is the number of covariates in the analysis, 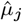 is the overall predicted probability of species presence at location *j, x*_*j*,*c*_ is the value of the covariate *c* in cell *j*, and 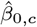 and 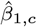 are the regression coefficients estimated in step 1 of the approach which characterize the relationship between the covariate *c* and the IK-informed probabilities of presence *Y*_*i*,*c*_.

If covariates with complex relationships (e.g., quadratic) are included we can estimate the final probabilities of species presence using

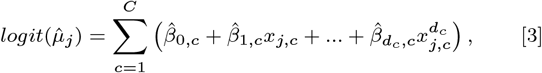

where *d*_*c*_ is the number of order for the polynomial for covariate *c*.

If time varying covariates are included we can estimate the final probabilities of species presence using,

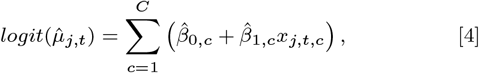

where *t* is time.

If all above scenarios are included in the analysis the final probabilities of species presence is estimated using,

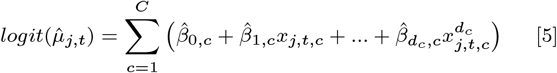

The prediction is mapped and then presented to the knowledge holders, if there is any noted discrepancy, we go back to step 1.

### Case Study Application

Indigenous Knowledge of ringed seal habitat use was documented as part of a collaborative research project with Iñupiat hunters in Utqiaġvik, Alaska (46)(Figure 1). At the beginning of the project, free prior and informed consent was obtained from the IK holders through a written and oral review of a summary of the project. The researchers and IK holders signed an agreement of how the IK would be documented, used, and shared. The agreement was revisited during subsequent interviews. All interviews conducted for this research were performed under the University of British Columbia Behavioural Research Ethics Board certificate H18-02585. Semi-directed interviews were conducted with nine active hunters focused on species-habitat associations and behaviour of ice-associated seals. After the initial interviews, the IK was summarized by habitat type and then reviewed by the hunters to ensure the summary reflected the IK shared. Detailed methods can be found in Gryba *et al*.(46).

We applied the methods to four important environmental covariates, identified by the hunters interviewed, that influence the habitat selection of ringed seals in summer in the waters around Utqiaġvik: distance from shore, ice concentration, currents, and spatial regions. Knowledge of distance from shore, a qualitative type of IK, was characterized as a simple linear relationship with a probability of 0.88 assigned to areas within 5 km of shore decreasing to probability of 0.33 120 km from shore (Figure 2A, Table 1). As outlined in Step 1, the probabilities were inferred from the proportion of hunters that shared IK of this species-habitat relationship. In this case, 8*/*9 hunters indicated association with nearshore habitat, and 3*/*9 hunters indicated association with offshore habitat. No transformation was applied as we were not predicting beyond the range of the data and the probabilities were within 0 and 1. Ice concentration was treated as a quantitative type of IK, and ringed seal presence was quantified for a set of ice concentration bins (i.e.,0%, 10-20%, 30-40%, 50-60%, 70-80%, 90%, 100%) (46). To reflect the increased probability of ringed seal presence in areas of high ice concentrations (i.e., July 2015, fig. S1), the probabilities were increased from 0.11 in ≥ 90% ice concentration (46) to 0.55 in 90% ice and 0.66 in 100% ice (Figure 2C). To characterize the relationship between the probability of ringed seal presence with ice concentration, we applied beta regression with a quadratic relationship to capture the higher probability of presence at low and high ice concentrations. The resulting estimate of probability of presence was mapped and reviewed by the IK holders (Figure 2D, E). The knowledge holders indicated that these increases better reflected their observations. Distance from shore and ice concentration were treated as continuous variables in the analysis, and scaled prior to modeling.

During the review of the probability of ringed seal presence maps, one hunter noted that the region of Admiralty Bay should have lower probability of presence (as in) (46). The probability of presence within the specified region was set to 0 and the areas outside had a probability value of 0.89 (Figure 4B, Table 1). To characterize this spatial IK, we created a categorical variable that equalled zero in the region specified by the IK and 0.89 in the area outside. Similarly, currents were analyzed as a spatial categorical variable: main current region, near the main current region, and far from the main area (outside current regions). Currents were considered a constant covariate as the IK indicated regions of high currents that are used by ringed seals throughout the summer. The locations of currents used by ringed seals in summer were based on areas indicated by hunters (46), a spatial IK. These areas were buffered to provide higher probability where the currents were identified and then decreasing probability 10-20 km from the main current areas (Figure 4D, Table 1). The region at the currents was given a probability of presence of 0.11 reflecting that each region was identified by one hunter (i.e., 1 hunter out of 9 hunters). The mapped probabilities were reviewed by the IK holders. The environmental variables used for mapping were projected to a 5 km x 5 km grid, Alaska Albers projection, with ice concentration from Spreen *et al*.(66).

The characterized relationships between the IK probability of presence and distance to shore, ice concentration, region, and areas of high currents were mapped for review. All analysis was completed in R (R Core Team 2021) and beta regression was completed using a Bayesian approach and the R package brms (67), with flat priors over the entire real line. Model convergence was checked using the Gelman-Rubin statistic and by visual inspection of chain plots. The beta regressions were run for 10,000 iterations with a warm-up of 2,000 iterations. The final estimate of the probability of ringed seal presence included all four covariates in equation 5. All examples of IK in this study and the resulting maps were reviewed by six of the IK holders who participated in the IK documentation conducted by R. Gryba *et al*.(46).

## Data, Materials, and Software Availability

The IK included in the case study is available in Gryba *et al*.(46). All R source code is available via GitHub….

## ACKNOWLEDGMENTS

Many thanks to the additional hunter who requested to remain anonymous for sharing their knowledge and reviewing all results. Many thanks to Taqulik Hepa for her thoughts, input, and continued support of this project. Thank you to F. Dupont and P. Molloy for reviewing and testing code and to T. Martin, F. Wiese, K. Chan, and M. Humphries for thoughtful comments and insights on this work. Additional thanks and credit to Danielle Larson of NorthWest Strategies for Figures 1, 2, and 3. This work was funded by the North Pacific Research Board (Project 1815), and supported in part by funding from the Social Sciences and Humanities Research Council Doctoral Fellowship, University of British Columbia Four Year Doctoral Fellowship, ACUNS Dr. Jim McDonald Scholarship, Lawrence Edward Hassell Graduate Field Research Award in Fisheries, Northern Scientific Training Program, Stantec Research & Development funds, Canadian Research Chairs program, Natural Sciences and Engineering Research Council of Canada, BC Knowledge Development fund, and Canada Foundation for Innovation’s John R. Evans Leaders Fund.

## Supplementary Materials

**Fig. S1.**
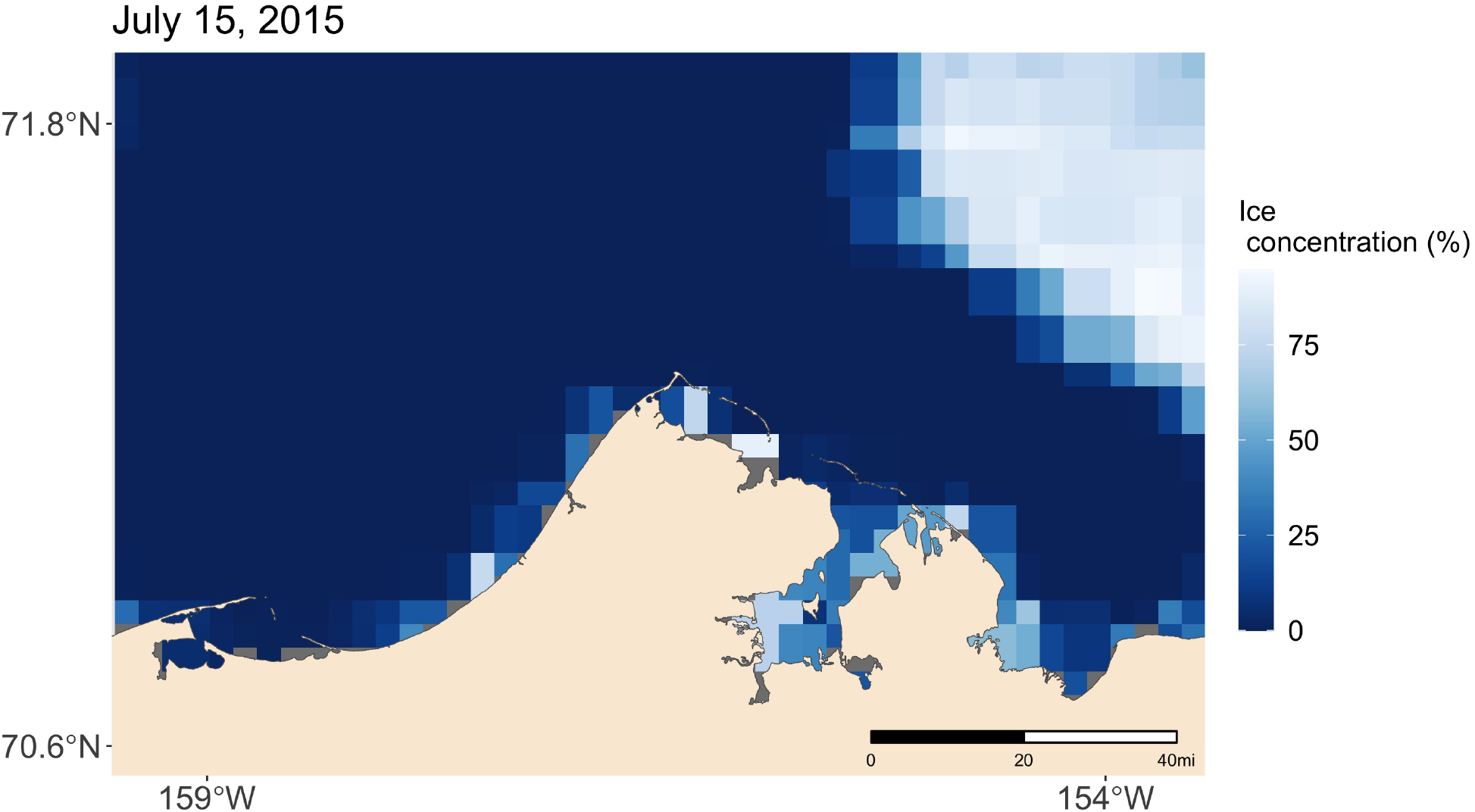
Ice concentration (%) in the area near Utqiaġvik, Alaska on July 15, 2015 (66). White is high ice concentration, dark blue is low ice concentration.

